# The efficacy and accuracy of ribosomal RNA depletion methods

**DOI:** 10.1101/2025.07.23.666324

**Authors:** Mohammed Ahmed, Charmaine Bishop, Andrea Betancourt, Vicky Hunt, Mark Viney

## Abstract

The depletion of ribosomal RNA (rRNA) is a critical step in RNA-sequence analyses, used to enhance the detection of non-rRNA molecules, such as messenger RNAs and non-coding RNAs. However, the efficiency and potential biases introduced by different rRNA depletion methods remain poorly characterized. Here, we evaluated three commercially available rRNA depletion kits – QIAseq FastSelect, riboPOOL, Zymo-Seq RiboFree – for their performance with the parasitic nematode *Strongyloides ratti*. We assessed the kits’ efficiency in rRNA removal, the recovery of expressed genes and transposable elements, and the detection of spliced leader sequences and genes’ operonic organization. Zymo-Seq demonstrated the highest sensitivity and minimal bias in a measure of gene expression, while QIAseq showed the least rRNA depletion and significant differential expression biases. Our findings underscore the importance of empirical validation of rRNA depletion methods, particularly for parasites and non-model organisms, and we suggest that Zymo-Seq as the optimal choice for *S. ratti* and related nematodes.

---

Cells contain a wide variety of types of RNA molecules including ribosomal RNA (rRNA), messenger RNA (mRNA), transfer RNA (tRNA), and small RNAs (sRNA). Among these, rRNA is the dominant type, constituting more than 85% of total cellular RNA in most organisms[1]. For many sequence-based studies, such as RNA-seq[2], it is desirable to eliminate or reduce the rRNA fraction, so to maximise the concentration of the non-rRNA molecules in the sample. For studies of mRNA, this is often done by poly-A selection[3]. For studies involving nematodes, the spliced leader (SL) sequence commonly added to the 5’ end of mRNAs[4] can also be used to select mRNA molecules[5]. For other studies, rRNA depletion is commonly used, which typically involves the hybridization of antisense DNA / RNA oligonucleotides to rRNA, and the subsequent removal of the resulting DNA / RNA-rRNA hybrid[3].

Most commercially available rRNA depletion kits are probe-based and work by specifically targeting rRNA with antisense oligonucleotides[6]. The main drawback to these methods is that they are sequence-specific, and so rely on sufficient specificity to the rRNA sequence of the target species to efficiently deplete rRNA, while not targeting other non-rRNA sequences. For species that are not commonly-used model species, species-specific probes may not be readily available. In contrast, the Zymo-Seq RiboFree method is a probe-free method, potentially widely applicable to most organisms. This method synthesizes complementary DNA (cDNA) from total RNA, deaturates then renatures RNA-DNA hybrids, then removes RNA-cDNA hybrids. Because of the abundance of rRNA, hybridization and subsequent removal of these molecules is efficient compared to other, less abundant, RNAs[7].

Despite the importance of rRNA depletion methods there have been few comparisons of the efficiency of different rRNA depletion methods or assessments of whether they introduce biases in RNA-seq data. Several studies have compared different rRNA depletion methods and / or poly-A selection methods[2,6,8]. Comparison of RiboZero rRNA depletion with poly-A RNA selection in seeking to identify long non-coding RNAs (lncRNAs) found that poly-A selection allowed detection of more lncRNAs, although rRNA depletion allowed better detection of other RNA classes[9]. A study optimizing RNA-seq library construction for low input RNA from *Caenorhabditis elegans* found that, compared with poly-A selection, rRNA depletion with custom-designed probes was effective by a number of measures[10]. A comparison of seven rRNA depletion kits concluded that all successfully removed a significant amount of rRNA, although the RNase H-based kits gave more consistent results compared to probe-based methods[11]. While rRNA depletion is capable of capturing both poly-A^+^ and poly-A^-^ RNA it has also been shown to introduce bias through overestimation of some types of RNA and capture of some immature mRNAs[2].

Here, we compare three commercial rRNA depletion kits (and a no rRNA-depletion control) for their suitability for use with the animal parasitic nematode, *Strongyloides ratti. S. ratti* is a widely recognised model for the study of parasitism, with substantial genomic resources[12– 14].

We made a single preparation of RNA from Trizol-stored *S. ratti* mixed free-living stages and from this used three different rRNA depletion treatment methods: (i) QIAseq FastSelect –rRNA Worm Kit for ribodepletion with NEBNext® Ultra™ II Directional RNA Library Preparation. This kit was designed for *C. elegans* but may work for other species depending on their rRNA sequence similarity to *C. elegans* (*S. ratti* identity to *C. elegans* 74-77% and 80-89 % for mitochondrial and nuclear rRNA, respectively); (ii) siTOOLs Biotech riboPOOL Depletion Kit for ribodepletion with NEBNext® Ultra™ II Directional RNA Library Preparation, where *S. ratti*-specific probes targeting both nuclear and mitochondrial rRNA (except the 5S rRNA) were designed by the manufacturer; (iii) Zymo-Seq RiboFree Total RNA Library Preparation, and (iv) a no rRNA-depletion control, which was done using a modified Zymo-Seq RiboFree Library Preparation protocol, but omitting the treatment of samples with depletion reagents. Each of the three treatments and the control were triplicated. Libraries were sequenced on the Illumina NovaSeq X Plus platform, and each library had at least 66 million reads post-trimming (**Supplementary Table 1**). Full experimental and bioinformatic methods are given in the Supplementary Information.

We then evaluated the three kits for (i) their efficiency in removing rRNA, (ii) the non-target removal or overestimation of expressed genes and transposable elements, (iii) how well they capture other RNA types and (iv) their utility in depicting trans-splicing and operonic organisation of genes through the analysis of the SL types associated with the transcribed genes.

We quantified the rRNA depletion by determination of the proportion of reads that mapped to rRNA. In the no rRNA-depletion control, rRNA-mapped reads were an average of 88.4 (SD 0.53) % of all reads, with 18S and 28S rRNA each representing an average of 40.8 (SD 1.72) and 43.5 (SD 1.23) % of the total reads, respectively (**Fig. 1**). All three rRNA depletion treatments reduced the proportional abundance of rRNA-mapped reads, but to varying extents. The QIAseq treatment resulted in the least rRNA depletion, only reducing rRNA-mapped reads to 68.9 (SD 0.22) %, with the depletion mainly of the 18S rRNA and to a very limited extent the two mitochondrial rRNAs. The riboPOOL treatment resulted in the greatest rRNA depletion, reducing rRNA-mapped reads to 20.4 (SD 5.78) %, with almost complete elimination of the 12S, 16S and the 5.8S mapped reads. The Zymo-Seq treatment was broadly comparable to the riboPOOL treatment (rRNA 41.37, SD 1.40) %, though less effective at reducing the 12S and 16S mitochondrial rRNA, inflating their proportional presence. In general, there was good agreement among the three replicates of each of the three treatments and the control.

**Fig. 1.**
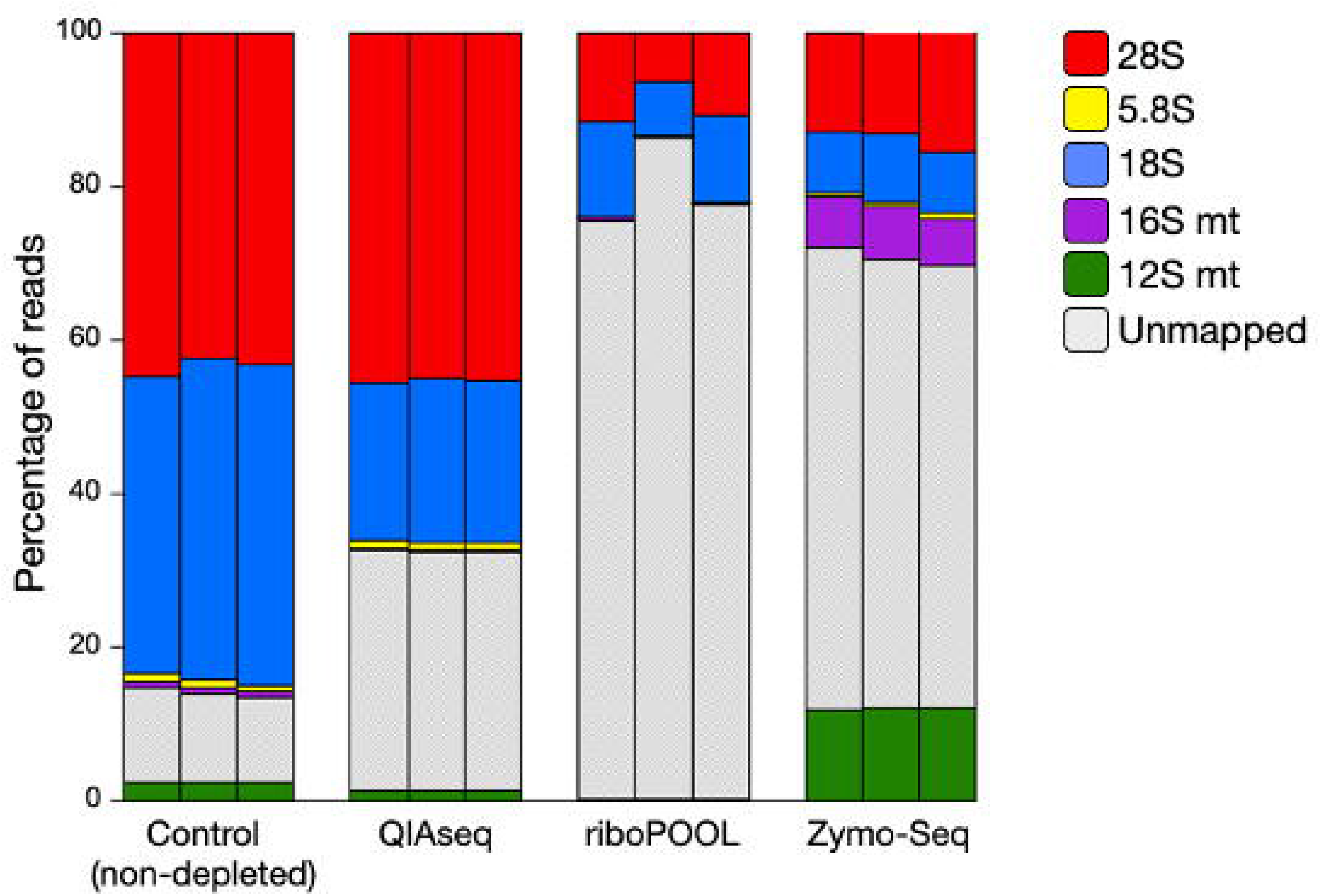
The proportional abundance of nuclear and mitochondrial rRNA in control RNA and following three rRNA depletion treatments, showing three replicates of each. Unmapped reads are non-rRNA reads.

We next compared the three depletion treatments and the control for the recovery of expressed genes using two metrics (i) sensitivity, the proportion of loci, transcripts, exons and introns that we could assemble and annotate from each library, and (ii) precision, the proportion of the recovered features that matched exactly with those in the *S. ratti* reference genome[15]. The results show that the Zymo-Seq treatment had the highest sensitivity for locus, transcripts, exons and introns, whereas the QIAseq treatment had the lowest for all four features (**Fig. 2A**). The QIAseq and riboPOOL treatments had lower sensitivities than the control. For precision, the three rRNA depletion treatments were very similar to the control for exon, introns and transcripts. For locus, the Zymo-Seq treatment had a higher precision than the control, and QIAseq and riboPOOL had a lower precision than the control (**Fig. 2B**).

**Fig. 2.**
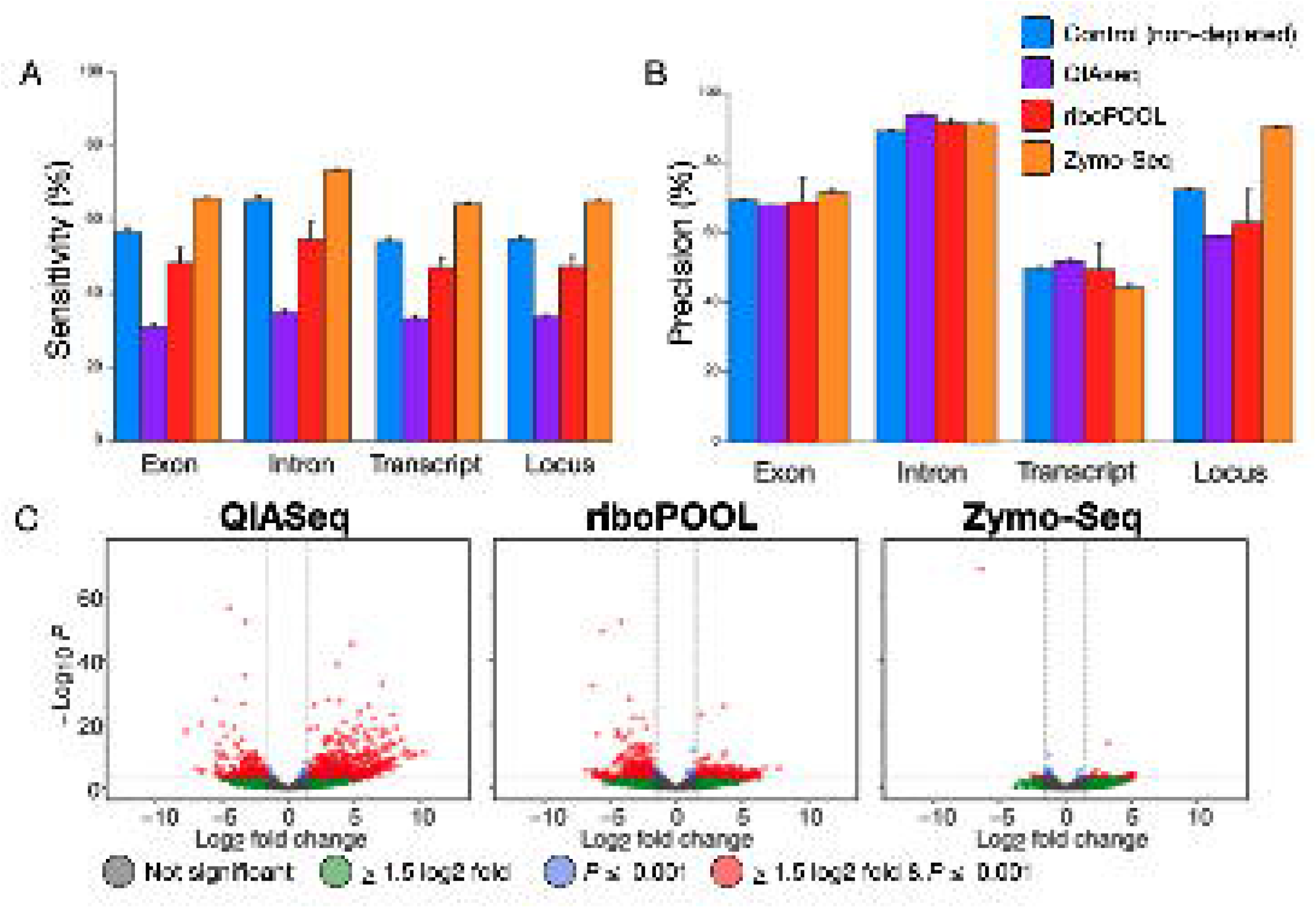
Recovery of expressed genes in control RNA and following three rRNA depletion treatments showing (A) Sensitivity (B) Precision. Values shown are averages of three replicates; error bars are + 1 SD. (C) Differential gene expression analysis. The log_2_ fold change is the difference in expression of the treatment from the no rRNA-depletion control. Each dot is one gene; colour coded as shown. Red are genes that are significantly differentially expressed (P ≤ 0.001) at ≥ 1.5 log_2_ fold change difference between the treatment and control; negative values have lower expression in the treatment compared to the control positive vice versa. Grey are genes with < 1.5 log_2_ fold change and not significant; green are genes with a > 1.5 log_2_ fold change but not significant; blue are genes < 1.5 log_2_ fold change but significant. Many dots overlie each other making the colours more intense. The y-axis is the negative log of P. The vertical dotted lines are 1.5 log_2_ fold change; the horizontal dotted lines are P = 0.001.

We also performed a differential expression analysis of genes between the no rRNA-depletion control and each of the rRNA depletion treatments, with any apparent differential expression between a library and the control potentially being due to a bias in the recovery of mRNA during rRNA depletion. Both QIAseq and riboPOOL resulted in the differential expression of 992 (700 upregulated; 292 downregulated) and 689 (368 upregulated; 321 downregulated) genes, respectively (**Fig. 2C**), with QIAseq libraries showing a bias towards overexpression (a positive log_2_ fold change), and riboPOOL almost equal overexpression and under-expression (a negative log_2_ fold change). Zymo-Seq showed the least differential expression (93 genes, 80 upregulated; 13 downregulated).

We investigated the presence of expressed TEs in the libraries and *de novo* annotated and classified a curated list of 38 TE families, and then compared the TE presence in the 3 treatments and the control. Overall, less than one percent of reads mapped to the TEs. The QIAseq and the control libraries had similar low rates of TE-mapped reads, whereas the riboPOOL and Zymo-Seq treatments had substantially higher rates (**Fig. 3A**). Long terminal repeat (LTR) retrotransposons were the most highly abundant TE in all treatments (**Fig. 3B**).

**Fig. 3.**
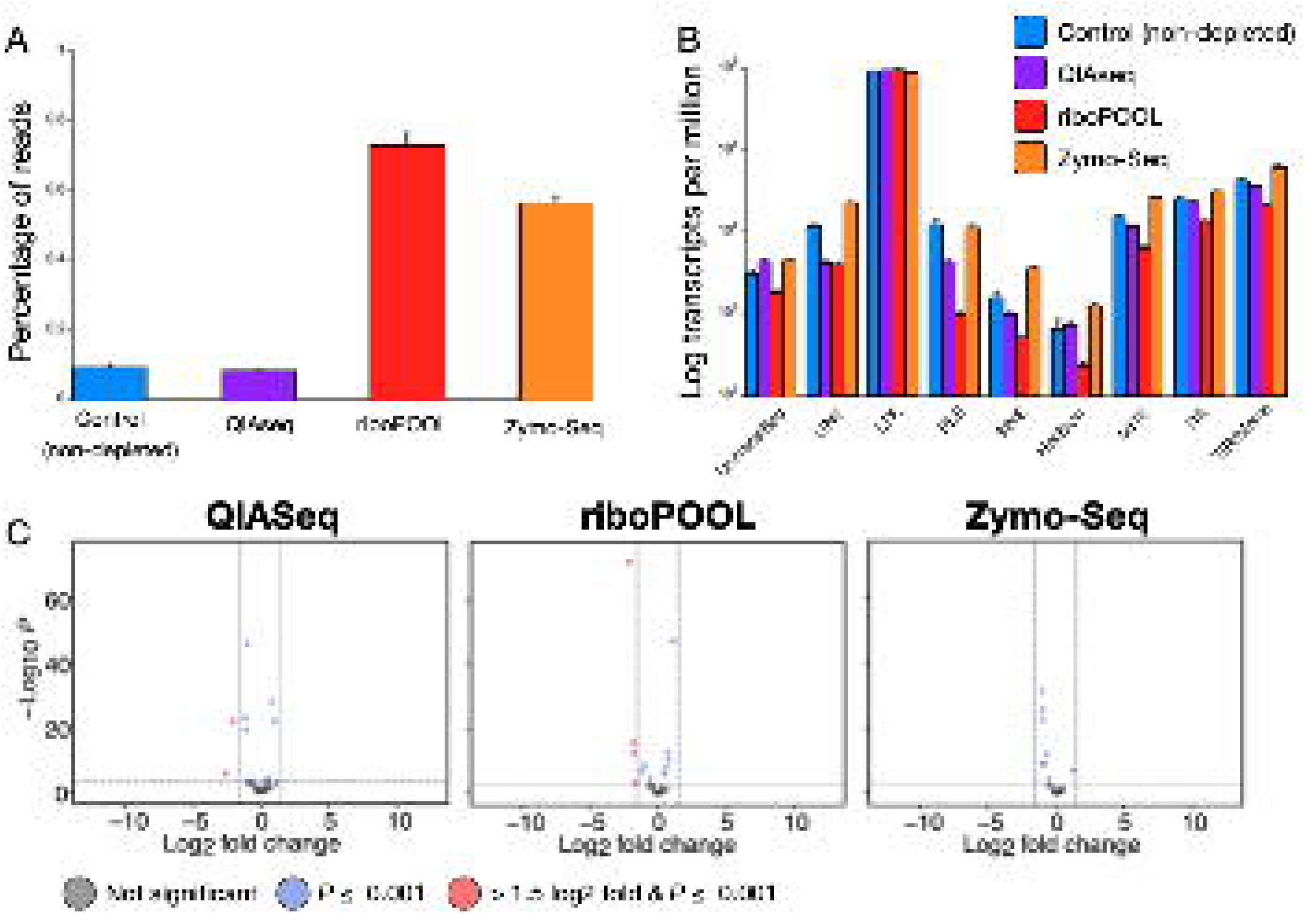
Expressed TEs (A) The proportional abundance of TE-mapped reads among the three treatments and the control; (B) the expression of TEs in log transcripts per million (TPM). Values shown are averages of three replicates; error bars are + 1 SD. (C) Differential TE expression analysis showing the log_2_ fold difference in expression of the treatment from the no rRNA-depletion control; negative values have lower expression in the treatment compared to the control; positive vice versa. Each dot represents one TE; colour coding as shown. Red are TEs that are significantly differentially expressed (P ≤ 0.001) at ≥ 1.5 log_2_ fold change difference between the treatment and control. Grey and blue are not significantly differentially expressed TEs. Grey are TEs with a < 1.5 log_2_ fold change and not significant; blue are TEs with a < 1.5 log_2_ fold change but significant. The y-axis is the negative log of P. The vertical dotted lines are 1.5 log_2_ fold change; the horizontal dotted lines are P = 0.001.

We investigated differential representation of TEs between each of the three treatments and the no rRNA-depletion control. The treatment that had the least difference from the control, which was Zymo-Seq, followed by QIAseq, and then riboPOOL. Both QIAseq and riboPOOL treatments showed decreased transcript abundance for some orders / superfamilies of TEs, while Zymo-Seq treatments showed no significant changes in TE expression (**Fig. 3C**).

We evaluated the efficiency of each treatment and control in detecting SL sequences and other non-coding RNAs. This identified an average of 28 (SD 0.6) consensus SL sequences from the three control, no rRNA-depletion libraries (**Table 1**; **Supplementary Table 2**). These formed two separate clusters: (i) an average of 17 (SD 4) belonging to a SL1-type cluster; (ii) an average of 11 (SD 4.2) belonging to a SL2-type cluster; and (iii) an average of <1 consensus sequences classified as being neither SL1 nor SL2 types. In the control libraries, an average of 5,035 (SD 67) genes (39.0% of all 12,908 genes in the reference genome annotation) were trans-spliced (**Table 1**). Most of them are monocistronic (average 4,549 (SD 57)), with only an average of 39 (SD 32.4) being operonic, organized into operons each consisting of exactly two genes.

**Table 1.**
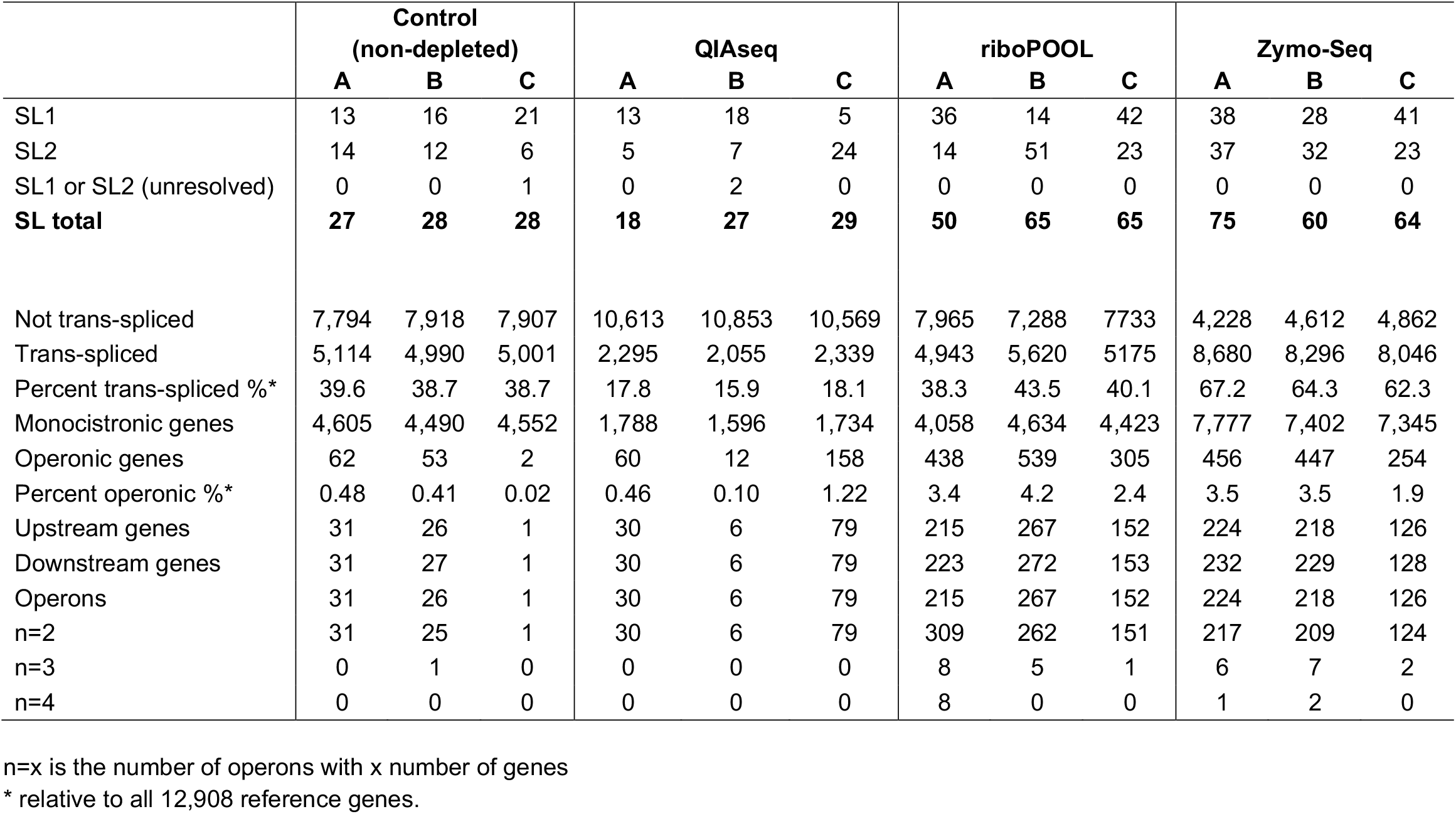
Number of SL sequences and their categorisations; trans-splicing events and operonic organization of genes in the control and the three rRNA depletion treatments as predicted using SLOPPR pipeline. Each SL represents a consensus SL sequences from several reads. For each treatment and control there are three replicates, A, B, C.

Of the three treatments only QIAseq, with an average of 25 (SD 5.9) SL sequences, resulted in the identification of fewer SL sequences than the control; riboPOOL and Zymo-Seq identified averages of 60 (SD 8.7) and 66 (SD 7.8) SL sequences, respectively. There was considerable variation among the treatments in the number of trans-spliced genes identified, ranging from an average of 17% (SD 1.2) (QIAseq) to 64.6% (SD 2.4) (Zymo-Seq) of all genes (**Table 1; Supplementary Tables 3 – 5**). Similarly, all three treatments resulted in the identification of different numbers of monocistronic and operonic genes. For all treatments and control, there were often considerable discrepancies between replicates in the number of SL sequences and the number of operonic genes identified. That said, the number of trans-spliced genes were consistent among replicates within treatments.

In this work we compared three different methods of rRNA depletion – Zymo-Seq, riboPOOL, and QiaSeq – on a single RNA extraction from the nematode *S. ratti*. Overall, there was wide variation in the levels of rRNA depletion, with QiaSeq resulting in minimal rRNA reduction. Further, there was variation in the inferred expression levels of genes due to two methods (riboPool and QiaSeq), suggesting mRNA recovery for these methods may not be unbiased. In contrast, apparent differential expression was minimal for libraries prepared using Zymo-Seq depletion. Finally, the Zymo-Seq treatment had the highest sensitivity overall, with the highest the recovery of expressed genes, expressed TEs, and SLs.

However, even in the no rRNA-depletion control, there was a less than 60% recovery of mRNA. This may be for two reasons, (i) many *S. ratti* genes will only be expressed in the within-host parasitic stages of the life cycle, and so RNA from these genes will not be present in the sample that we have analysed, and / or (ii) because of genes with a very low level of expression at, or beyond, our limit of detection. Previous expressed sequence tag analysis of different *S. ratti* life cycle stages showed that 38% (of 4,152 sequence clusters) were expressed in only the within-host parasitic female stages [16]. This is consistent with the recovery of 60% of expressed in genes in the present study from free-living stages.

Our study has limitations. Firstly, our no rRNA-depletion control library was prepared using the Zymo-Seq RiboFree Library preparation method, which is a different library preparation method than used in the two other rRNA depletion treatments, so that we cannot formally differentiate the effect of rRNA depletion method and library preparation method, however, our experimental design does test and compare combined, rRNA depletion and library preparation methods. Two, our study only analyses *S. ratti*, and it would be desirable to extend these comparisons to other parasitic nematodes, and to other parasites more generally.

We conclude that the Zymo-Seq rRNA depletion method is the best of the tested methods for *S. ratti*, as it achieves a substantial rRNA depletion while generally having the least effect on the representation of the remaining RNA molecules. Our results further suggest that it may be important to experimentally test different rRNA depletion methods for specific purposes and organisms.

## CRediT author contribution statement

**Mark Viney**: Conceptualisation, Funding acquisition, writing– original draft, review and editing. **Vicky Hunt**: Conceptualisation, Funding acquisition, writing– review and editing. **Andrea Betancourt:** Funding acquisition, writing– review and editing. **Mohammed Ahmed**: Conceptualisation, writing– original draft, data analysis. **Charmaine Bishop**: Resources– production of free-living *S. ratti*.

## Acknowledgments

We would like to thank the Centre for Genomic Research, University of Liverpool for their help with this work and for providing access to computational resources used in this research. This work was supported by a grant from the BBSRC, BB/X008673/1.

