## Supplementary information for "The efficacy and accuracy of ribosomal RNA depletion methods"

**Supplementary Methods**

**Nematode material.** We collected approximately 20,000 *S. ratti* mixed free-living stages (infective larvae and free-living males and females) from faecal cultures of *S. ratti*-infected rats, which were transferred to 200 µL of Trizol (Thermo Fisher Scientific) and stored at -70ºC.

**RNA isolation.** All libraries were constructed from a single extraction of total RNA. To extract RNA, the Trizol-frozen nematodes were thawed, transferred to Biomasher tubes (Polysciences, Germany) on ice, and homogenised in three cycles of freezing on dry ice followed by disruption using the pestle on wet ice. The resulting homogenate was supplemented with cold Trizol to a volume of 1 mL, mixed by pipetting and incubated at room temperature for 5 minutes to allow for complete dissociation, after which 200 µL of 1-bromo-3-chloropropane was added, mixed by shaking, incubated for 3 minutes, then centrifuged for 15 minutes at 12,000 x g at 4ºC. The upper aqueous phase (containing the RNA) was transferred to a fresh tube to which 500 µL isopropanol, 5 µg of glycogen and 100 µL of 3 M sodium acetate was added, and the mixture held at -20ºC for 1 hour, after which it was centrifuged for 10 minutes at 12,000 x g at 4ºC, the supernatant discarded and the pellet washed in 70 v/v % ethanol, then centrifuged at 9,000 x g for 5 minutes at 4ºC, and the supernatant discarded; this was repeated three times. All remaining traces of ethanol were removed after the final centrifugation and the pellet was finally suspended in RNase-free water and incubated at 55ºC for 15 minutes.

**rRNA depletion and library preparation.** From the single RNA sample, three different rRNA depletion treatments and library preparations were carried out: (i) QIAseq FastSelect –rRNA Worm Kit for ribodepletion with NEBNext® Ultra™ II Directional RNA Library Preparation. This kit was designed for *C. elegans* but may work for other species depending on their rRNA sequence similarity to *C. elegans* (*S. ratti* identity to *C. elegans* 74-77% and 80-89 % for mitochondrial and nuclear rRNA, respectively); (ii) siTOOLs Biotech riboPOOL Depletion Kit for ribodepletion with NEBNext® Ultra™ II Directional RNA Library Preparation; (iii) Zymo-Seq RiboFree Total RNA Library Preparation, and (iv) a no rRNA-depletion control. Each of the three treatments and the control were triplicated. All libraries were stranded.

The QIAseq FastSelect -rRNA and NEBNext® Ultra™ II Directional RNA Library Preparation rRNA depletion and library preparation was carried out following the manufacturers’ instructions starting with 100 ng of total RNA for each replicate. Fragmentation time was 15 minutes. The rRNA-depleted RNA was then used as input material for the NEBNext Ultra II Directional RNA library prep kit using twenty-five-fold diluted adaptor. After 13 cycles of amplification, the libraries were purified using AMPure XP beads.

The siTOOLs riboPOOL Depletion and NEBNext® Ultra™ II Directional RNA Library Preparation required the design of *S.* *ratti* specific probes targeting both nuclear and mitochondrial rRNA, except the 5S rRNA. We submitted these sequences (**Supplementary Data 1**) to siTOOLs Biotech who designed the probes. Ribosomal RNA depletion and library preparation were performed using the siTOOLs Biotech riboPOOL Depletion Kit for ribodepletion, followed by NEBNext® Ultra™ II Directional RNA Library Preparation kit, both following the manufacturers’ instructions, with 200 ng of total RNA for each replicate. We cleaned the depleted RNA using CleanMag Beads (Paragon Genomics), then used 5 µL as the input material for the NEBNext Ultra II Directional RNA library preparation kit for Illumina as for the QIAseq method.

The Zymo-Seq RiboFree library preparation was performed using the Zymo-Seq Ribofree ® Total RNA Library Kit (Zymo Research) following the manufacturer’s instructions, with 100 ng of total RNA for each replicate, and 2 µL of cDNA synthesis reagent as the input material. Incubation time was 60 minutes and 11 cycles of amplification were performed.

The no rRNA-depletion control was processed using a modified Zymo-Seq RiboFree Library Preparation protocol (above), but omitting the treatment of samples with Depletion Reagents.

**Sequencing.** We quantified each of the 12 libraries (three treatments and control, with three replicates of each) after processing using a Bioanalyzer and by qPCR using the Illumina Library Quantification Kit following the manufacturer's instructions, diluted the DNA to 180 pM, and sequenced the libraries on the Illumina NovaSeq X Plus platform (Illumina®, San Diego, USA), generating 2 x 150 bp paired-end reads. We excluded reads < 15 bp long; reads were trimmed to remove adapters, low-quality bases (Q < 20) and all flanking “N” bases from reads using cutadapt [1].

**Assessment of rRNA depletion efficiency.** The rRNA depletion was quantified bioinformatically using FastQ screen v0.15.3[2]. To do this, for each replicate of the three treatments and the no-depletion control, sequence reads were mapped to a database of both nuclear and mitochondrial rRNA sequences using bowtie2[3] inside FastQ-screen (using default parameters)[2]; we did this separately for forward and reverse reads, and then took the average of the proportions, so giving a single value of unmapped reads for each library. We also recorded the proportion of reads mapping to each sequence in the database. The rRNA database consisted of the same full-length sequences of the 18S rRNA, 28S rRNA, 5.8S rRNA, 12S and 16S mitochondrial rRNA of *S. ratti* used to design the riboPOOL custom probes.

**Recovery of expressed genes.** We compared the three depletion treatments and the control for their recovery of expressed genes by calculating the proportion of the features: (i) locus, (ii) transcripts, (iii) exons and (iv) introns (as defined in the GffCompare[4] documentation, https://ccb.jhu.edu/software/stringtie/gffcompare.shtml) in the *S. ratti* annotated reference genome[5] (WormBase version WS289). To do this we used the rRNA-filtered reads (above), mapped them to the *S. ratti* reference genome[5] using STAR v2.7.8a[6] (--outSAMtype BAM Unsorted --runThreadN 32 --outSAMstrandField intronMotif), then sorted the alignments by coordinates using samtools v1.18[7]. We then generated transcript assemblies for the alignments with StringTie v2.2.3[8] (default parameters) using the reference genome and annotation as guide. The identification of coding regions within the StringTie transcripts was performed using Transdecoder v5.7.1[9] (default parameters). Finally, GffCompare v0.12.6 (default parameters) was used to determine the proportion of loci, transcripts, exons and introns present in the reference annotation that were recovered in each of the 12 libraries separately.

We also investigated the differential expression of genes between each depletion treatment compared to the no rRNA-depletion control. To do this we used the sorted read alignments generated during the StringTie annotation pipeline and then used featureCount[10] to count reads mapped to genes (-p -t transcript -g gene_id -F ‘GTF’ –primary -s 1). The read counts were then used in DESeq2[11] for differential gene expression analysis. Plots were generated using ggplot2 inside the R statistical package.

**Recovery of expressed TEs.** The quantity of expressed TEs recovered from each of the 3 treatments and the control was assessed using SalmonTE v0.4[12]. To do this we used a curated library of TEs and the raw reads from the 12 libraries together as the input for the SalmonTE in the quant mode (--exprtype=TPM). The curated TE library was made by first performing *de novo* TE classification using Earl Grey[13], then further curated using MCHelper[14], which performs automatic curation of TE libraries to reduce the incidence of false positives, reduce redundancy and improve TE classification. No manual curation was performed on the resulting library after MCHelper. The three treatments and control were also compared by the level of expression of different TE superfamilies.

We investigated the differential expression of TEs between the depletion treatments and the no rRNA-depletion control using DESeq2 (as above, for expressed genes) and then SalmonTE’s test mode (--tabletype=csv --analysis_type=DE). Plots were generated using ggplot2 in R.

**Recovery of other RNA molecules.** SL are commonly added to the 5’ end of nematode mRNA molecule[15,16]. We evaluated the efficiency of each treatment and control in detecting SL sequences and other non-coding RNAs. To do this we used a combination of the two pipelines, SLIDR and SLOPPR[17] to *de novo* identify SLs and to predict operons, respectively. SLIDR was run to identify all consensus SL sequences and their corresponding genes by supplying the reference genome and annotation[5] (WormBase version WS289) as inputs together with the rRNA-filtered reads of each of the treatments. The predicted SL sequences were then used as inputs for SLOPPR to determine how the SL sequences are organised into SL1 and SL2. SLOPPR produces clusters representing SL1 and SL2 types. We identified SL1 as that which was more frequently trans-spliced to monocistronic genes and to the upstream gene in the operons, which is the pattern found in *C. elegans[15]*. The resolved SL1-SL2 organisation was also used by the SLOPPR pipeline to determine the trans-splicing events as well as the operonic organisation of genes[17].

**Data availability**

All raw sequence data were deposited in the NCBI Sequence Read Archive (SRA) under the BioProject accession number PRJNA1243282 (SRR32905192–SRR32905203) at <https://www.ncbi.nlm.nih.gov/sra?term=SRP573992>.

**References to Supplementary Methods**

[9]	BJ. Haas, TransDecoder, (n.d.).

[15]	T. Blumenthal, Trans-splicing and operons in C. elegans, WormBook: The Online Review of C. Elegans Biology [Internet] (2018).

[16]	E.L. Lasda, T. Blumenthal, Trans‐splicing, Wiley Interdiscip Rev RNA 2 (2011) 417–434.

[17]	M.A. Wenzel, B. Müller, J. Pettitt, SLIDR and SLOPPR: flexible identification of spliced leader trans-splicing and prediction of eukaryotic operons from RNA-Seq data, BMC Bioinformatics 22 (2021) 140. https://doi.org/10.1186/s12859-021-04009-7.

**Supplementary Table 1.** Number of reads for each library, before and after trimming. A, B and C are the three replicates of each treatment and the control.

| **Sample** | **Raw Reads** | **Trimmed reads** | **rRNA-filtered reads** |
| --- | --- | --- | --- |
| Control A | 78,348,896 | 78,270,091 | 8,426,206 |
| Control B | 66,655,365 | 66,582,244 | 6,721,017 |
| Control C | 76,474,639 | 76,393,999 | 7,376,216 |
| QIAseq A | 163,003,286 | 162,760,013 | 49,387,566 |
| QIAseq B | 149,873,983 | 149,626,465 | 44,766,141 |
| QIAseq C | 149,764,403 | 149,521,589 | 44,904,725 |
| riboPOOL A | 161,682,269 | 161,361,518 | 120,933,644 |
| riboPOOL B | 213,546,394 | 213,163,227 | 183,265,609 |
| riboPOOL C | 207,124,911 | 206,697,950 | 159,152,974 |
| Zymo-Seq A | 145,674,583 | 144,880,325 | 85,797,115 |
| Zymo-Seq B | 117,332,474 | 116,708,572 | 66,476,511 |
| Zymo-Seq C | 136,532,819 | 136,003,525 | 76,525,238 |

**Supplementary Tables 2-5 are Excel sheets.**

**Supplementary Data 1 is a text file.**
